# Long-term evolution of *Streptococcus mitis* and *Streptococcus pneumoniae* leads to higher genetic diversity within rather than between human populations

**DOI:** 10.1101/2023.11.22.568241

**Authors:** Charlotte Davison, Sam Tallman, Megan de Ste-Croix, Martin Antonio, Marco R. Oggioni, Brenda Kwambana-Adams, Fabian Freund, Sandra Beleza

## Abstract

Evaluation of the apportionment of genetic diversity of bacterial commensals within and between populations is an important step in the characterization of their evolutionary potential. Recent studies showed a correlation between the genomic diversity of human commensal strains and that of their host, but the strength of this correlation and of the geographic structure among populations is a matter of debate. Here, we studied the genomic diversity and evolution of the phylogenetically related oronasopharingeal healthy-carriage *Streptococcus mitis* and *Streptococcus pneumoniae*, whose lifestyles range from stricter commensalism to high pathogenic potential. A total of 119 *S. mitis* genomes showed higher within- and among-host variation than 810 *S. pneumoniae* genomes in European, East Asian and African populations. Summary statistics of the site-frequency spectrum for synonymous and non-synonymous variation and ABC modelling showed this difference to be due to higher historical population effective size (*N_e_*) in *S. mitis,* whose genomic variation been maintained close to mutation-drift equilibrium across (at least many) generations, whereas *S. pneumoniae* has been expanding from a smaller ancestral population and has been subjected to adaptive selection. Strikingly, both species show limited differentiation among populations. As genetic differentiation is inversely proportional to the product of effective population size and migration rate (*N*_e_*m),* we argue that large *N_e_* have led to similar differentiation patterns, even if *m* is very low for *S. mitis*. We conclude that more diversity within than among human populations and limited population differentiation must be common features of the human microbiome due to large *N_e_*.

## Introduction

There is evidence that healthy carriage of human-associated bacteria have coevolved with humans over thousands of years [1–3], and that this has generated significant genetic diversity among hosts [4]. Reconstructions of within-species single nucleotide variants (SNVs) from metagenomic samples from the gut showed that bacterial strain diversity in human populations is globally consistent with theoretical expectations of long-term evolution by stochastic fluctuation of allele frequencies over the generations (genetic drift) and purifying selection[4, 5]. However, these studies have also uncovered a wide range of genetic variation among species. This is reflective of independent evolutionary histories enabling bacteria with different ecological trajectories. Importantly, these evolutionary histories have resulted from interactions with humans that range from bacteria with a stricter commensal lifestyle to others exhibiting high pathogenic potential. Characterization of the genetic variation of human-associated bacterial species on different ends of the pathogenic potential spectrum and the evolutionary mechanisms leading to that variation is, therefore, crucial for understanding bacterial adaptation to their host environments, as well as the evolution of phenotypes such as virulence.

An interesting case of phylogenetic related species that show contrasting interactions with humans are the two upper respiratory tract inhabitants *Streptococcus mitis* and *Streptococcus pneumoniae*. *S. mitis* is one of the most prevalent species of the oropharyngeal microbiome [6, 7], where it resides as a commensal. Because of its abundance at birth and throughout life, *S. mitis* is an excellent model for comprehensive analysis of variation and diversification of the oral microbiome. Compositional similarity between mother and child indicates vertical (familial) transmission of the oral microbiome [8]. However, early investigations into genetic diversity of *S. mitis*, although impeded by difficulties in distinguishing related species within the Mitis phylogenetic group [3, 9], found multiple genotypes in new-born infants both matching and not matching parental genotypes, suggesting some degree of horizontal transmission of the Mitis group including *S. mitis* [10, 11]. High *S. mitis* diversity within and between hosts is corroborated by analyses of the highly differentiating gdh gene, of a combination of multi-locus sequence alignment (MLSA) genes, and of a small number of genomes [12–14]. These patterns were associated both with relaxed immune pressure exerted by the host due to *S. mitis* commensal lifestyle, and with their primarily vertical mode of transmission leading to lineage isolation and, consequently, to preclusion of potential homogenization of the mitis gene pool due to homologous recombination [13, 14]. According with this view, discrete clusters of genetic variation tracking host genomic variation across populations should be observed, for instance in a similar manner to the stomach-colonizer and vertically transmitted *Helicobacter pylori* [15]. Recent studies have shown this to be the case for gut microbiome species [16], although the correlation between host and commensal genetic diversities seems to be weak [17–19]. The apportionment of genetic diversity of human commensals within and between human populations is still a subject of debate, and the factors that contribute to population differentiation need further consideration. This is important because the amount of population variation within and among populations contribute to the evolutionary potential of a species.

On the other hand, *S. pneumoniae* is a frequent colonizer of the nasopharynx that is associated with high human mortality and morbidity worldwide [20]. Although nasopharyngeal colonization is usually asymptomatic, it is considered an essential step preceding invasive and non-invasive pneumococcal disease [21, 22]. In addition, carriage serves as a source of pneumococci that can be transmitted to other people. This property of horizontal transmission will influence the level and distribution of genetic variation in the overall population of *S. pneumoniae* across host populations and, therefore, pneumococcal carriage is a key stage in the evolution of this organism. However, *S. pneumoniae* population genetic variation has been mostly characterised in relation to invasive disease [23–31] or in the context of vaccination epidemiological transition [32–40], and few recent studies address the genetic variation of carriage *S. pneumoniae* within a population [41–46].

A dominant mode of vertical transmission in *S. mitis* versus a dominant mode horizontal transmission in carriage *S. pneumoniae* is expected to lead to more geographic isolation of the former in comparison to the latter, and this is expected to lead to the observation of higher geographic structure in *S. mitis* than in *S. pneumoniae*. However, under the standard neutral model, genetic differentiation among populations is inversely proportional to the product of effective population size and migration rate (*N*_e_*m;* [47]). This means that in populations with large *N_e_*, even in cases where m is minimal, effective migration will overcome genetic drift (proportional to 1/*N_e_*) leading to low long-term population differentiation.

Here, we sought to gain a better understanding about the evolution and population differentiation of healthy carriage of *S. mitis* and *S. pneumoniae* across human populations. We collected bacterial samples from European, East Asian and African hosts, which compose the major divisions of human genetic diversity [48], in this way, sampling across deeper evolutionary time scales. We observe that the apportionment of genetic diversity is higher within populations than between populations, and that this is related with the large *N_e_* for both species. However, genomic variation is considerable higher for *S. mitis* due to a bigger ancestral population that has been maintained close to mutation-drift equilibrium across (at least many) generations; in *S. pneumoniae,* contemporary diversity is inferred to be due to a population expansion from an ancestral population with smaller *N_e_* and the action of adaptive selection.

## Results

### *S. mitis* has considerably higher within-host diversity than carriage *S. pneumoniae*

We collected a representation of within- and between-host genomic variation of healthy colonisation of *S. mitis* and *S. pneumoniae* across major human population groups. The *S. mitis* dataset consists of a total of 101 newly collected and 18 publicly available isolates from 46 independent hosts from Africa (n= 17 hosts/32 isolates), East Asia (n= 6 hosts/37 isolates), and Europe (n= 24 hosts/50 isolates). More than one isolate was collected from 18 hosts, ranging from 2 to 10 isolates per host. The African dataset included 20 isolates from five pairs of related individuals (two mother-child pairs, two sibling pairs and one sibling trio), whereas the European and East Asian isolates were obtained from unrelated individuals only.

To obtain a sample of the true general diversity of carriage *S. pneumoniae* populations that is not biased by the action of strong recent selection, the *S. pneumoniae* dataset was obtained from published pre-vaccination studies [39, 41, 43] and comprises a total of 810 isolates from asymptomatic hosts (carriage isolates) from Africa (n= 90 hosts /230 isolates), East Asia (n= 480 hosts/isolates) and Europe (n= 100 hosts/isolates). The African dataset included within-host genomic variation, with the number of isolates collected from 71 hosts ranging from two to five.

Genetic diversity and presence of clonal relationships within the host, within families and between hosts were assessed for both species by calculating the number of single nucleotide variant (SNV) differences between every pair of isolates in the total dataset and within each of those three levels of host relatedness. For *S. mitis*, the mean number of pairwise SNV differences in the total sample is 30,190 (Interquartile range, IQR: 29,050 - 32,580). We observed clonal relationships in ∼one percent of total pairwise comparisons (<1000 SNVs), mostly from within-the host, but also between related individuals (two sibling pairs) and between unrelated individuals’ isolates collected in the same geographic region (five pairs; Supplementary Fig. 1A). Pairwise SNV comparisons within the host (mean= 24,366; IQR: 25,298-30,358) and between related hosts (mean= 25,411; IQR: 24,822-32,769) are slightly smaller than pairwise SNV comparisons between unrelated hosts (mean=30,531; IQR: 29,189-38,379) (Kruskal-Wallis test, p-value≤7.08×10^-5^; Supplementary Fig. 1C).

In *S. pneumoniae,* the presence of clonal relationships was detected both within and between hosts at a larger degree than in *S. mitis* (Supplementary Fig. 1B and Supplementary Fig. 1D). The mean number of pairwise SNV differences between serotypes is 7,309 (IQR:6,310-7,748), similar to the mean number of pairwise SNV differences within serotypes (mean=7,265, IQR: 5,348-10,481). Considering genomic variation in carriage *S. pneumoniae* organised in Global Pneumococcal Sequence Clusters (GPSCs) of shared evolutionary history as previously defined [49, 50], the mean number of pairwise SNV differences is greater between (7,393; IQR: 6,319-7,866) than within the defined clusters (1,880, IQR: 128 – 3,113). Analysing the within-the-host pairwise SNV differences distribution in the African sample, we observe that ∼97% of the pairwise comparisons within GPSCs and within serotypes are smaller than 100; the remaining pairwise SNV comparisons are above the first quartile of between-host pairwise SNV differences. We then set a threshold of 100 pairwise SNV differences to define clonal relationships in *S. pneumoniae*.

Considering the established cut-off for clonality for both species, we observe higher within-host diversity for *S. mitis*: 64 out of 242 (26%) and 155 out of 236 (66%) within-host pairwise comparisons are defined as clonal relationships for *S. mitis* and *S. pneumoniae*, respectively. We can conclude that human hosts are generally colonised by multiple and divergent strains of *S. mitis*, whereas *S. pneumoniae* populations within the host are most often dominated by a single strain.

As we are interested in investigating global population genetic structure and long-term evolutionary dynamics of both species, we extracted the maximal datasets composed solely by unrelated strains. In total, 75 *S. mitis* isolates (25 African, 32 European, 18 Asian) from 45 hosts, and 353 *S. pneumoniae* isolates (78 African, 68 European, 207 Asian) from 339 hosts from across the three geographic regions were used in the analyses.

### *S. mitis* has considerably higher within-population between-host diversity than carriage *S. pneumoniae*

Phylogenetic analyses of geographically restricted (mainly European) *S. mitis* and mainly invasive *S. pneumoniae* genomes have shown higher diversification of *S. mitis* than *S. pneumoniae* [14]. Here, we analyse if these patterns of genome variation for both species are observed in our extended dataset of exclusively carriage isolates from across geographically distant regions, which corresponds to analysing the scale of diversity in these species associated with host ancestry and with deeper evolutionary timescales.

We first evaluated the sets of genes available to both species for their evolutionary success. The pangenome – defined as the total set of genes observed across all sampled isolates - and mean individual genome sizes are similar across the three geographic regions for *S. mitis* and *S. pneumoniae* (Supplementary Table 1). To compare the pangenome and genome sizes between the two species, we have analysed 1000 random samples of unrelated *S. pneumoniae* with the same size as *S. mitis* (n=75) and present the mean estimates. Individual genome size is smaller for *S. mitis* – on average 1832 genes (SD=84) for *S. mitis* and 2005 genes (SD=71) for *S. pneumoniae* (Mann-Whitney test, *P*-value< 2.20 x10^-16^). However, *S. mitis* gene repertoire is significantly larger than that of *S. pneumoniae*. In a total of 75 strains, the *S. mitis* pangenome was estimated to harbour 9626 genes, of which ∼10% were found in all strains (core genes, 951 genes). At the same level of resolution, *S. pneumoniae* pangenome size is 6064 genes (mean across 1000 random samples; SD=250), of which 18% comprise core genes (mean ± SD=1092 ± 29). Furthermore, the impact of each additional genome on the size of the pan genome is greater for *S. mitis* than *S. pneumoniae* (Fig. 1). Analysing permuted data with power law regression [51], we estimated that each *S. mitis* genome adds on average 2.8 times more genes to the pangenome than any *S. pneumoniae* genome (Fig. 1).

**Figure 1.**
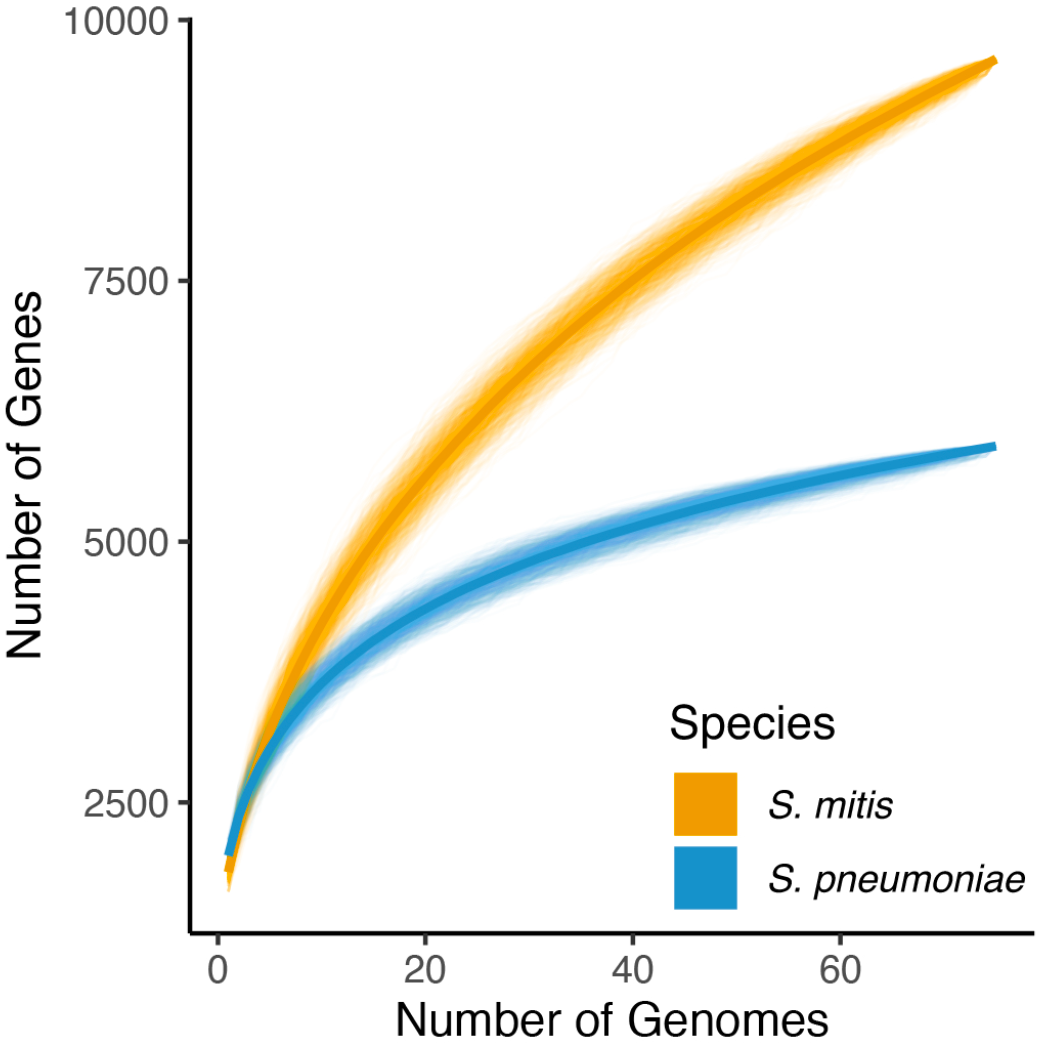
*S. mitis* and *S. pneumoniae* pangenomes according to power law fit. The number of genes is plotted as a function of the number of genomes. In *S. mitis*, the order of the 75 unrelated genomes was permuted 1000 times. In *S. pneumoniae*, the same procedure was applied to each of 1000 random samples of 75 unrelated genomes. Power law regression was fitted to the mean number of genes obtained across all permutations. Fitting the same model to the median gave similar results, albeit with lower goodness-of-fit to the *S. pneumoniae* data. The parameterisation that best fitted the data was: Y = *a*X*^b^* + *c*. For *S. mitis*, *b* = 0.42; for *S. pneumoniae*, *b* = 0.12 – 0.32; mean=0.2. *S. mitis*, orange; *S. pneumoniae*, blue.

Based on the core genomes extracted for each species, nucleotide diversity (π) within species, estimated accounting for core genome length and sample size, was similar across the three geographic locations: for *S. mitis*, π values for the core genome are 0.034, 0.032 and 0.033 for the African, Asian and European populations of isolates, respectively; for *S. pneumoniae*, π values for the core genome are 0.008, 0.010 and 0.008 for the African, Asian and European populations of isolates, respectively. Considering the species as a whole, π is higher in *S. mitis* (π = 0.034) than in *S. pneumoniae* (π=0.010). *S. mitis* is therefore considered to have substantially higher genetic diversity than its’ close relative pneumococcus in terms of both pangenome content and sequence variation.

### Limited population differentiation in both *S. mitis* and carriage *S. pneumoniae*

Phylogenetic analysis and PCA show both species to have little genetic differentiation among geographic locations (Fig. 2; Supplementary Fig. 2). Concordant with these analyses, between population divergence values in *S. mitis* and *S. pneumoniae* among geographic regions are >3-fold lower (average Hudson’s F_ST_= 0.043 in *S. mitis*; 0.039 in *S. pneumoniae*; Supplementary Table 2) than that in humans from the same geographic regions (average Hudson’s F_ST_= 0.135; Bathia et al. 2013). Analysis of molecular variance confirms that most of the genetic variation in *S. mitis* and *S. pneumoniae* is segregating within rather than between geographically distinct populations: the within-population component explains 95.3% and 96.3%, whereas the among-geographic-regions component explains 4.7% and 3.7% of the genetic variation, respectively for *S. mitis* and *S. pneumoniae*.

**Figure 2.**
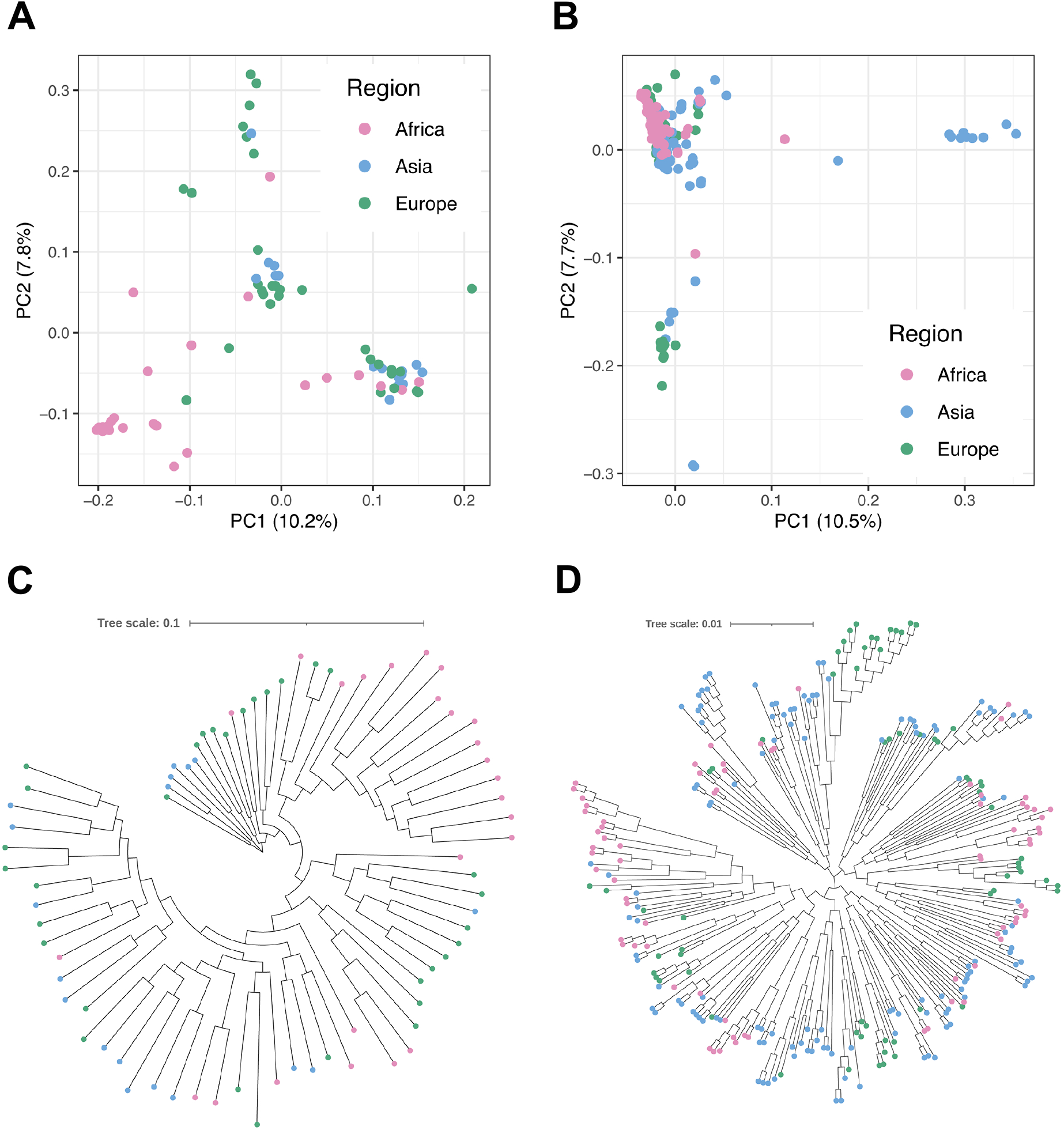
Population genetic structure in *S. mitis* and *S. pneumoniae*. **A.** plots of PC1 versus PC2 for *S. mitis*. **B.** Plots of PC1 versus PC2 for *S. pneumoniae*. **C.** Maximum likelihood unrooted phylogenetic tree for *S. mitis.* **D.** Maximum likelihood unrooted phylogenetic tree for *S. pneumoniae*. Tree scales are given in substitutions per core genome site. Analyses presented here for *S. pneumoniae* do not include outlying serotype NT isolates, whose higher genetic variation has been attributed to higher recombination rates for this clade [41]. Graphical representation of the analyses for the full *S. pneumoniae* sample are in Supplementary Fig. 2. Colour code corresponds to geographic region: Africa, pink Asia, blue; Europe, green.

We can conclude that, although the two species conform to contrasting population models of transmission and dispersal, both *S. mitis* and *S. pneumoniae* genomic variation is mostly randomly assorted according to geography and host ancestry. The lack of strong population differentiation in both species means that the evolutionary history of the isolates can be modelled by assuming that isolates form a single, non-structured population.

### Long-term evolutionary dynamics for *S. mitis* and carriage *S. pneumoniae*

The considerably higher level of *S. mitis* genetic diversity in comparison to *S. pneumoniae* can be attributed to deterministic (mutation and recombination) and population level processes (genetic drift and selection) which dictate the frequency of the allelic variation in the population. We first determined that effective mutation (the product of *N_e_* and mutation rate) and effective recombination (the product of *N_e_* and recombination rate) are not responsible for genetic diversity differences between the two species (Supplementary Note 1).

We then computed the minor-allele frequency distribution of variant sites, called the folded site-frequency spectrum (_f_SFS; Fig. 3A and Fig. 3B), independently for synonymous and nonsynonymous variation, each to evaluate the role of genetic drift (derived from demographic history of the species) and selection occurring during each species’ evolutionary history, respectively [52]. Notably, we inferred high recombination rates for both species (Supplementary Note 1), and therefore, we assume that selection only affects a reduced number of closely linked neutral sites around the selected variant.

**Figure 3.**
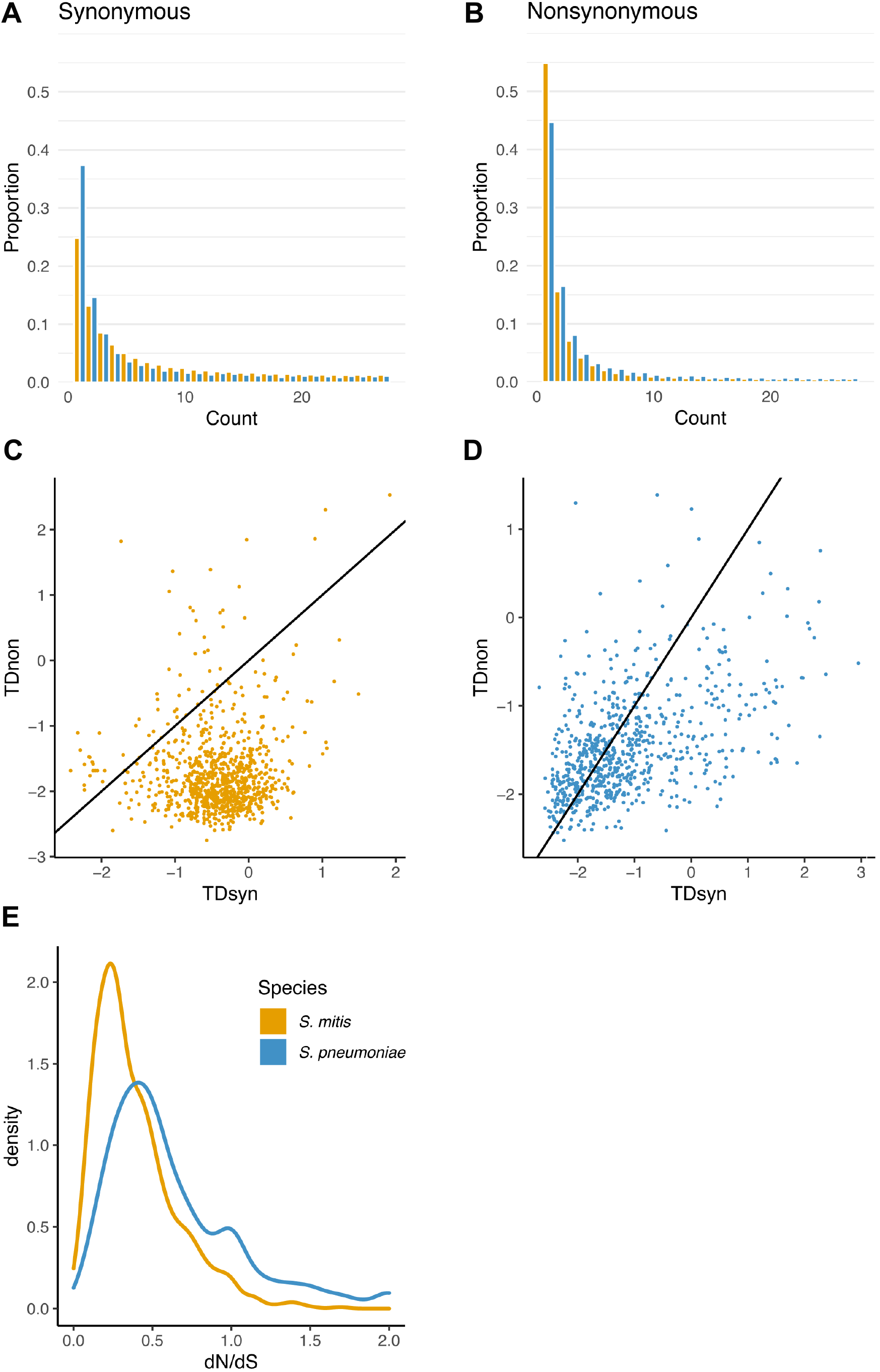
Evolutionary dynamics of *S. mitis* and *S. pneumoniae.* **A** Folded site frequency spectrum for synonymous variation. **B** Folded site frequency spectrum for nonsynonymous variation species. To get a comparable result, the SFS was calculated based on the same sample size for both species. **C** Tajima’s D for synonymous variation (TDsyn) vs Tajima’s D for nonsynonymous variation (TDnon) in *S. mitis*. Line corresponds to the diagonal (where x=y). **D** Tajima’s D for synonymous variation (TDsyn) vs Tajima’s D for nonsynonymous variation (TDnon) in *S. pneumoniae.* Line corresponds to the diagonal (where x=y). **E** d*N*/d*S* distribution for both species, truncated at d*N*/d*S*=2. *S. mitis*, orange; *S. pneumoniae*, blue.

*S. pneumoniae* shows an excess of rare synonymous (<2%) variants compared to *S. mitis* (chi-square (1df), p-value < 2.20 x10^-16^) (Fig. 3A). In contrast, a higher number of intermediate frequency synonymous variants (>5%) is observed in *S. mitis*. Conversely, the overall *S. mitis* nonsynonymous _f_SFS is skewed towards rare variation in comparison to *S. pneumoniae* (chi-square (1 df), p-value < 2.20 x10^-16^; Fig. 3B). The distributions of synonymous variation suggest contrasting demographic models of non-growth versus population growth in *S. mitis* versus *S. pneumoniae* and, potentially, different types and magnitudes of selection.

To test for these hypotheses, both species’ _f_SFS were summarised by calculating Tajima’s D (TD) per core genome and per core gene, for synonymous (TDsyn) and nonsynonymous (TDnon) variants (Fig. 3C and D). Overall core genome TDs were compared with simulated TDs obtained under a standard population equilibrium and selective neutral model (Methods). In *S. mitis*, TDnon is consistently smaller than TDsyn (Fig. 3C). The overall TDsyn is −0.36 (p-value= 0.380), whereas TDnon is −1.91 (p-value = 0.003). This confirms that the evolutionary model most consistent with *S. mitis* genomic variation is one of a stationary population in equilibrium (at least over many generations) and natural selection acting at non-synonymous sites. In contrast, there is a linear tendency for larger values of TDnon to be associated with larger values of TDsyn in *S. pneumoniae*, and both overall metrics are significantly smaller than zero (overall TDnon = −1.90, p-value=0.001; overall TDsyn= −1.40, p-value=0.039). This confirms that the _f_SFS of *S. pneumoniae* is more consistent with a model of population growth influencing the scale of genetic diversity observed in *S. pneumoniae* and natural selection acting at nonsynonymous site*s*. Approximate Bayesian computation (ABC) based on a random forest classifier of ∼25,000 simulations of 100kb windows obtained under an exponential growth model which incorporated recombination and mutation parameters estimated for *S. pneumoniae* shows support for significant population growth (growth rate of 15.64; controlling for regions under strong selection) in the *S. pneumoniae* population (Supplementary Note 2).

The ratio of the number of nonsynonymous substitutions to the number of synonymous substitutions (d*N*/d*S*) per base pair (calculated per gene and normalised for gene length; see methods) allows us to further investigate the role of natural selection in *S. mitis* and *S. pneumoniae* populations (Fig. 3E). Whereas *S. mitis* d*N*/d*S* distribution tends towards lower d*N*/d*S* values (mean= 0.228; median= 0.157; IQR: 0.097-0.268), evidencing a preponderance of purifying selection acting at nonsynonymous sites, the *S. pneumoniae* d*N*/d*S* distribution is shifted to larger values (mean= 0.745; median= 0.514; IQR: 0.341-0.889) and has an extended right-hand tail, evidencing the action of both purifying and adaptive selection acting at nonsynonymous sites.

### Long-term evolution of effective population sizes

Neutral genetic diversity within populations is determined by the population effective size and the mutation rate, a notion that is captured in the population parameter *θ*. As we confirmed experimentally that the mutation rate and growth cycle are comparable between the two species (Supplementary Note 3), differences in *θ* between *S. mitis* and *S. pneumoniae* will be due to their *N_e_*. We have then inferred the coalescent *N_e_* using the observed number of segregating sites for each species (which ≈ *θE*(*L_n_*), where *E*(*L_n_*) is the expected length [in coalescent time units] of the genealogy of the sample - Methods; we excluded the regions under selection identified in Supplementary Data Files 1-4 in the calculation of *S_n_*), and controlling for the population expansion in *S. pneumoniae* (Methods). Considering the mutation rate from [32], this indicated that *S. mitis*’s contemporary *N_e_* is only 1.5x higher than *S. pneumoniae*’s *N_e_*: *S. mitis N_e_*= 249,210*; S. pneumoniae N_e_*= 165,965 ([150,028–176,286] calculated based on the median, and minimum and maximum growth rates - Supplementary Note 2). We conclude that the ancestral population of *S. pneumoniae* prior to expansion was significantly smaller than the concurrent *S. mitis* population and that *S. pneumoniae*’s recovery to similar contemporary *N_e_* was not sufficient to generate comparable genetic diversity levels to *S. mitis*.

### Long-term adaptive evolution in *S. mitis* and *S. pneumoniae*

To investigate the putative action of selection at a deeper evolutionary scale, we compared the functions of the genes in the 5^th^ (generally considered under purifying selection) and 95^th^ (generally considered under adaptive selection) percentiles of the d*N*/d*S* distribution (Supplementary Data Files 1-4).

There was an over-representation of genes annotated as being involved in “cell wall/membrane/envelope biogenesis” and “defense” in the upper bound of both *S. mitis* and *S. pneumoniae* d*N*/d*S* distributions (Fisher exact test, p-value< 0.05). In *S. mitis*, all these genes are involved in self-immunity against bacteria and other perturbations in their host environment, such as antibiotic intake. Namely, SM12261_RS01820 encodes for an intramembrane metalloprotease with a domain that shows 96% identity to CAAX proteases involved in self-immunity against bacteriocins in *S. pneumoniae* (the targets of these proteases are not known; [53, 54]. *pbp1a* and SM12261_RS05940 encode for the penicillin-binding protein 1a and a serine hydrolase with 95% identity to domains of the beta-lactamase protein family in *Streptococcus* sp., both which are involved in bacterial resistance to beta-lactams [55–57].

In *S. pneumoniae*, the list of genes in the abovementioned categories includes *galE,* which encodes for an epimerase that is essential in the biosynthesis of polysaccharide capsule and of O-antigen polysaccharide units [58, 59], molecules whose main function is to protect against complement-mediated and phagocytosis killing [60–62]; SPR_RS03695 encoding for hemolysin, which was shown to play a crucial role in regulation of the capsule expression and in anti-phagocytosis [63]; SPR_RS09480 encoding for ribonuclease E containing a LysM peptidoglycan-binding domain and *ybeY* encoding for a ribonuclease Y, both which are involved in mRNA decay and in regulating gene expression during changes of the host environment, such as exposure to macrophages [64, 65]. Another category of genes includes *licD1,* which encodes for a transferase that mediates the incorporation of choline residues in peptidoglycan teichoic and lipoteichoic acids, which facilitate adhesion to the host [66–68]. Finally, *vanZ* is associated with tolerance to lipoglycopeptide antibiotics such as teicoplanin and vancomycin, commonly used in clinics [69]. In summary, genes under adaptive selection in *S. pneumoniae* are associated with self-immunity, colonization and modulation of host-inflammatory response.

## Discussion

Here, we have studied the genomic diversity, population differentiation and evolution of two phylogenetically related human-associated bacterial species, whose modes of transmission, dispersal, and interaction with host lead to different expectations in the magnitude of genomic variation (higher for *S. mitis*) and of population differentiation/structure (more significant for *S. mitis*). Indeed, we have determined higher historical *Ne* in *S. mitis* that has led to more substantial levels of genomic diversity in this species compared to *S. pneumoniae*, even though contemporary *Ne* between species is more comparable. However, both species present limited geographic differentiation across host populations contrary to our expectations based on modes of transmission. We now discuss how different patterns of natural colonization, transmission, dispersal, interactions with the host, and demographic histories have led to these populational patterns.

Contrary to *S. pneumoniae (*our data; [44]) and common species in the gut microbiome [5, 70], where a dominant strain is retained over prolonged periods of the host life, our results show that *S. mi*tis exhibits multiple colonization. According to [12] diversifying lineages may coexist for long stretches of time, generating within-host variation that will be able to respond to strong selection pressures such as changes in diet or antibiotic intake. The presence of multiple highly divergent lineages within the same host makes *S. mitis* less amenable to be studied using metagenomic samples; here we developed a culture system that allows to obtain isolates for within-the host studies more effectively. Our data confirms that additional *S. mitis* lineages are acquired within households but can also be acquired horizontally within contained social networks, as recently described for the oral microbiome more generally [71, 72]. Multiple colonization and some degree of horizontal transmission leads to effective recombination between strains (Supplementary Note 1), contrary to what was proposed [13].

We replicate the previously observed higher levels of genome variation in *S. mitis* than in *S. pneumoniae* [13] and extend these observations to the pangenome level. We determined that *S. mitis* variation is consistent with the most common evolutionary model for human commensals of population in mutation-drift equilibrium, with some degree of purifying selection [4, 5]. The higher *Ne* calculated for this species leads to a highly diverse populational gene pool (Supplementary Note 1) within which horizontal gene transfer (HGT) can occur. We additionally confirm that *S. mitis* pangenomes are shaped by cross-species HGT at a higher rate than *S. pneumoniae pangenomes* (only 79% of the reads in the *S. mitis* dataset are identified as streptococcal as opposed to 93% of the reads in the *S. pneumoniae* dataset using KrakenUniq [73]; Welch’s t-test p-value <0.0001). Although the *S. mitis* pangenome diversity can be explained by neutral evolution [74], future work should indicate if common accessory genes confer adaptive advantage within and across populations [75].

Modelling of *S. pneumoniae* genomic diversity agrees with a population expansion as a demographic model for *S. pneumoniae*. As *S. mitis* and *S. pneumoniae* have the same host, the distinct demographic history between both species must be linked with behavioural differences. As [14] proposes, the property of horizontal transmission makes *S. pneumoniae* more dependent on having enough hosts for successful dispersal and, consequently, population growth in S*. pneumoniae* is most likely intimately linked with the increase in human population density [76]. In addition, we argue that the horizontal transmission features of continuous founding bottlenecks and recovery are analogous to range expansion processes, and these properties can lead to the same genomic patterns as an (instantaneous) population expansion if the number of ‘migrants’ or bacterial cells between hosts is large (*Nm*>>1, where *N* is the within-host population effective size; [77]). Within-host deep-sequencing showed transmission bottlenecks sizes between donor and recipient to be higher than one [44], the within-host diversity of one colonization event (defined from cultured isolates) is consistent with *N_e_* ranging from 1 to 72 bacterial cells [43], and mice colonization experiments allowed to estimate within-host *N_e_s* of ∼100 [78]. In fact, a multiple merger coalescent genealogy was previously inferred for *S. pneumoniae* [79], which is concordant with range expansion [80]. It would be interesting to understand how horizontal transmission affects *Ne* over a great number of generations. Another aspect of *S. pneumoniae*’s genetic diversity is that it is also due to higher signals of adaptation (Fig. 3E; Supplementary Note 2), which can generate dynamics similar to recurrent bottlenecks and contribute to reducing long-term *N_e_* [81].

An important question that has been debated recently is the amount of commensal strain variation within and among different geographic regions or ethnic groups [16, 17]. Here we show that two important components of the oronasopharyngeal microbiome in humans show little differentiation across human populations. In an island model under neutral equilibrium, *F_ST_* is equal to 1/(1+2*N_e_m*) in haploids where *m* is the migration rate [47]. Studies on the transmissibility of the oral genome showed that strain dispersal across populations is minimal but not null [72]. Therefore, for *S. mitis* the large *N_e_* leads to values of *N_e_m* >>1, even if *m* is very small, reducing the differentiation among populations. In contrast, there are reports of pathogenic outbreaks extending for long geographic distance within Europe and across the world for *S. pneumoniae* [32, 82], and we detected pairs of highly related strains between continents (within our dataset 350 pairs of strains involving 125 strains from different geographic regions had <2000 SNP pairwise differences [<1%percentile]), demonstrating meaningful migration rates. Therefore, although ancestral *N_e_* is smaller in *S. pneumoniae*, it may have been and currently is sufficiently high enough that together with higher *m* make *N_e_m* >>1. In conclusion, our results support the view that more diversity within than among populations and little population differentiation must be common features of the human microbiome due to large *N_e_*.

## Material and Methods

### Bacterial Isolate collection

African *S. mitis* samples were collected by MRC The Gambia Unit, under ethical approval granted by the MRC-Gambia Ethics committee. Buccal swabs are part of an extended dataset collected from across The Gambia from family trios consisting of mother, child (3-10 years old) and a baby (less than 2 years old). Suspected streptococcal species (based on plate morphology and presence of alpha-haemolysis) were analysed via MALDI-TOF mass spectrometry, and putative Mitis group isolates had their genomes sequenced (see below). Mitis group species identification was done analysing MLSA variation as in [83]. The final dataset consisted of 32 isolates from one baby/two siblings’ trio, two sibling/baby duos, two mother/child duos, and six unrelated individuals.

European and East Asian samples were collected under ethics approval granted by Ethics Committee for Medicine and Biological Sciences at the University of Leicester (protocol 14610). All samples were collected in Leicester, UK. European samples were collected from the cheeks and tongue individuals natural and living in the UK for the past 2-5 years. East Asian samples were collected from five Chinese individuals that had lived in the UK for less than six months, maintained a Chinese diet and had either no romantic partner or a Chinese partner who met the same criteria. We complemented these datasets with publicly available European and East Asian whole genomes, which we confirmed via phylogenetic and F_ST_ analyses that they did not differ significantly from our two sample sets.

*S. pneumoniae* carriage samples matching the broad geographic origin of the *S. mitis* dataset (Europe: UK, East Asia: Thailand and Africa: The Gambia]) were collated from the literature [39, 41, 43]. Serotypes and sequence types are disclosed in respective publications and genomic sequences are available on the European Nucleotide Archive under study accessions: PRJEB2357 (Asian dataset), PRJEB2417 (European dataset), and PRJEB3084 (African dataset). We only analysed isolates from non-vaccinated individuals. For the East Asian dataset, we considered the subset of isolates collected in the first year of a 3-year sampling period, to obtain a random sample of a size comparable to the other studied geographic regions. This subsample might include more than one isolate per individual, but because we did not have access to that information, we did not consider it for within-host analyses. For the African and European datasets, we considered only the within-host carriage isolates that were collected pre-vaccination.

### *S. mitis* isolation and growth

The European and East Asian samples were cultured on Mitis Salivarius Agar (5% Sucrose (Fisher Scientific), 1.5% Peptone (Oxoid), 1.5% agar (BioGene), 0.5% Tryptone (Oxoid), 0.4% di-potassium hydrogen orthophosphate (Fisons), 0.1% Glucose (Fisons), 7.5E-3% Trypan Blue (Sigma), 8E-5% Crystal Violet (Sigma))[84]. Putative *S. mitis* colonies were identified based on flat, “rough” morphology and a blue colouring in the centre of the colony following 18 hours growth at 37°C, 5% CO2, and on the sequencing of the house keeping gene *glucose-6-dehydrogenase* (*gdh*) for unambiguous differentiation between Mitis group species [12].

This isolation technique had a significantly better yield than that used for African samples: 95% of isolates predicted as *S. mitis* based on *gdh* phylogeny clustering were confirmed as *S. mitis* based on whole genome data (European and East Asian isolates) compared to 37% of isolates where species was predicted with MALDI-TOF mass spectrometry (African isolates).

### DNA extraction, sequencing and *de novo* sequence assembly

DNA extraction was performed using Wizard Genomic DNA purification kit (Promega) and manufacturer’s instructions for Gram-positive bacteria. Samples were sequenced on an Illumina HiSeq 4000 at the Oxford Genomics Centre (UK), to a mean coverage of 83.4x for African isolates and 300x for East Asian and European isolates.

Raw FASTQ reads were quality normalised with Trimmomatic 2.0 [85]. Quality was confirmed via FastQC 0.11.5 [86]. Trimmed, paired reads were assembled to contig level using SPAdes 3.12 [87]. Assembly was improved to scaffold level using the “assembly improvement” pipeline from Sanger Pathogens [88]. Assembly quality was assessed via QUAST 4.3 [89]. The mean N50 across isolates was 239.6Kb (42.3Kb-931Kb) and the GC% content (39.2%-40.6%, mean=40.1%) was mirrored across isolates. Accession numbers for this dataset are in Supplementary Table 4.

### Serotyping and determination of Global Pneumococcal Sequence Clusters

Capsular serotyping and determination of Global Pneumococcal Sequence Clusters (GPSCs) of shared descent for *S. pneumoniae* [49, 50] were performed by uploading newly assembled genomes in the web tool PathogenWatch (https://pathogen.watch/). Two genomes from the Asian *S. pneumoniae* dataset were classified as *S. pseudopneumoniae* and were therefore removed from the dataset.

### Core and accessory genome extraction and pangenome analysis

Scaffolds were annotated using Prokka 1.14.6 with default settings [90] and the full set of non-redundant genes was extracted with Roary 3.13 [91]. Alignment was made invoking MAFFT [92] and considering a sequence identity of 85%. The percentage sequence identity was empirically determined, by considering the percentage that minimised the change in the number of genes per change in the parameter. Core genomes were formed as a concatenation of genes identified to be present in 100% of input sequences. The total core genome sizes obtained were 900,605 bp for *S. mitis* and 816,449bp for *S. pneumoniae*.

To evaluate differences of pangenome size between *S mitis* and *S. pneumoniae*, we performed an iterative procedure in which we permuted the input order of the bacterial genomes and assessed the rate of new genes identified per addition of each genome. For *S. mitis*, the input order of the (unrelated) genomes was permuted 1000 times. For *S. pneumoniae*, because final sample size of (unrelated) isolates was significantly larger than for *S. mitis*, we employed the same procedure to each of 1000 random samples of unrelated genomes of the same size as the one for *S. mitis*. Obtained curves were fitted applying power law regression to the mean number of number of genes obtained across permutations [51].

### Phylogenetic analysis

Phylogenetic trees were built from FASTA alignments with FastTree 2.1 [93], using the generalised time-reversible model of nucleotide evolution and re-scaled based on likelihoods reported under the discrete gamma model with 20 rate categories. This is a standard approximation for accounting for variable evolutionary rates across sites and uncertainty in these rates [94]. Trees were visualised using both iTOL [95].

### Variant and gene mapping and annotation

Trimmed fastq files were mapped to type strains *S. mitis* NCTC 12261 (NCBI accession: NZ_CP028414.1) and *S. pneumoniae* R6 (NCBI accession: NC_003098.1) using BWA, and variants called using SAMtools 1.3.2 mpileup [96, 97]. In order to be called a core variant, a site read depth of >10% of the mean genome-wide read depth, as well as minimum sequencing and mapping quality scores of 30, were implemented.

Synonymous and nonsynonymous variants were called in comparison to the type strains using snpEff [98]. To generate genetic diversity estimates per gene (see below), we considered the gene coordinates recorded in the genome assemblies of type strains.

### Genetic diversity, population structure, and folded Site Frequency Spectrum analyses

These analyses used biallelic variant sites only. Pairwise SNV differences were calculated with the software SNP-dists 0.6 [99]. Pairwise F_ST_ between populations and nucleotide diversity (π) of core genomes were calculated with the python package scikit-allel [100]. We calculated Hudson’s F_ST_ estimator which is less influenced by differences in sample size and SNV ascertainment scheme [101]. Pairwise F_ST_’s are based on the set of SNVS segregating in both populations; however, considering the set of SNVs segregating in either population gave similar F_ST_ values. Analysis of Molecular Variance (AMOVA) based on the pairwise differences between sequences was performed in ARLEQUIN ver3.5.2.2 [102].

Principal component analyses (PCA) of population structure were conducted in PLINK v1.9 [103]. PLINK V1.9 was also used to determine the Minor allele frequency (MAF) of variant sites, which was used to generate the _f_SFS for synonymous and nonsynonymous sites.

### Per Gene analyses

Nucleotide diversity (π), Waterson’s θ, and Tajima’s D (TD) per gene and for synonymous and non-synonymous substitutions independently using scikit-allel. We tested if the TD statistic deviated from a population equilibrium and selective neutral model by generating 10,000 random samples under this model and with the same diversity as the one observed for both species in ARLEQUIN ver3.5.2.2 [102]; p-values of the D statistic correspond to the proportion of random D statistics less or equal to the observation. If this fraction is <0.05, we concluded that the observed TD is significantly different from zero and not evolving according to the standard neutral model. We then used observed TDsyn statistics to determine the action of specific demographic processes (TDsyn<0 is expected under population growth, and TDsyn>0 under population contraction), and observed TDnon statistics to determine the action of natural selection (TDnon< 0 is expected under purifying and positive selection and TDnon>0 are expected under balancing selection) on the species’ genetic variation.

To assess the distribution of non-synonymous (d*N*) to synonymous (d*S*) variation at core genes, we computed the ratio of Waterson’s θ for non-synonymous variants to the Waterson’s θ for synonymous variants, normalised by gene length, to obtain a d*N*/d*S* per bp. EggNOG (v5.0) database was used for functional annotation based on Orthologous Groups (OGs) of proteins at different taxonomic levels [104].

### *N_e_* calculation

*N_e_* and magnitude difference in *N_e_* between species was calculated using coalescence theory and the observed number of segregating sites, *S_n_*. Ignoring the effect of selection, a sample of size n from a haploid species is expected to have E(*S_n_*) ≈ *S_n_* = N_e_*mu*E(L_n_), where E(L_n_) is the expected length (in coalescent time units) of the genealogy of the sample [105], and *mu* is the per-genome, per-generation mutation rate (*mu*=*μgl*, where *μ* is the pneumococcal mutation rate of 1.57 × 10^−6^ site^−1^year^−1^ [32], *g* is the generation time of 14/cell divisions /year [43], and *l is* the core genome size for *S. mitis* and *S. pneumoniae*). We have then inferred *N_e_* considering a constant size model for *S. mitis*, where 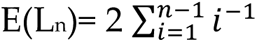 [105], and a population expansion from *S. pneumoniae,* where E(L_n_) was calculated using the recursion from [106], as implemented in [107].

## Acknowledgements

This work was supported by MRC-UK MR/M01987X/1 awarded to SB; by BBSRC-UK and University of Leicester funded Midlands Integrative Biosciences Training Partnership (MIBTP), grant number BB/M01116X/1, to CD; University of Leicester Doctoral Program Training Fellowship to ST. We thank Ebony Cave for bioinformatic support during the writing of this paper.

## Competing Interests

The authors declare no competing interests.

## Data Availability Statement

The *S. mitis* genomic dataset was deposited in NCBI-SRA (https://www.ncbi.nlm.nih.gov/sra) and are available under accession numbers provided in Supplementary Table 4. *S. pneumoniae* genomes were collected from the European Nucleotide Archive (ENA, https://www.ebi.ac.uk/ena/browser/home), available under the study accession numbers: PRJEB2357 (Asian dataset), PRJEB2417 (European dataset), and PRJEB3084 (African dataset). The reference genomes sequence used for the read mapping are available from GenBank (*S. mitis* NCTC 12261, NCBI accession: NZ_CP028414.1, GenBank accession: GCA_000960005.1; and *S. pneumoniae* R6, NCBI accession: NC_003098.1, GenBank accession: AE007317.1).

## Notes

### Competing Interest Statement

The authors have declared no competing interest.

