## Supplementary material for "Long-term evolution of *Streptococcus mitis* and *Streptococcus pneumoniae* leads to higher genetic diversity within rather than between human populations": Davison_etal_Supplementary material_biorxiv.pdf

#### Supplementary Notes

##### Supplementary Note 1. Assessing the role of mutation and recombination in generating genetic diversity in *S. mitis* and *S. pneumoniae* populations

Recombination and mutation population parameters were estimated using the coalescence analytical framework implemented in mcorr [1] (Supplementary Table 3), which is appropriate for species composed of highly divergent strains and for which the underlying phylogeny is difficult to estimate accurately. In addition, the estimation of evolutionary parameters in mcorr is based on synonymous variation only, which minimises the effect of selection. We first performed a gene-by-gene alignment of core genes using Prokka and Roary outputs. Resulting fasta files were converted to eXtended Multi-FAsta (XMFA) format [2], which were used as input files in mcorr. Only sequences with <2% gaps of the total alignment length were used in the analysis. mcorr computes the correlation profile, P(l), in each species by averaging over the correlation profiles of synonymous substitutions for all gene sequence pairs in the sample, and then averaging over all genes. When homologous recombination is present, the probability of observing a correlated substitution (P(l)) decreases with distance between any two loci (l), at a rate that is proportional to the recombination rate. Otherwise, the function is constant. By fitting P(l) to an analytical form inferred based on a coalescence model with recombination, mcorr estimates parameters that characterize the more diverse species' gene pool with which the sampled genomes have recombined: the mutational divergence ($\theta_{pool}=2\overline{T}\mu$, where $\mu$ is the mutation rate), the recombinational divergence ($\phi_{pool}= 2\overline{T}\delta$, where $\delta$ is recombination rate), the relative rate of recombination to mutation ($\theta_{pool}/\phi_{pool}= \delta/\mu$), and the mean recombination fragment length ($\overline{f}$), where $\overline{T}$ is the mean pairwise coalescence time across all loci in the overall population ($\overline{T}=N_e/2$) in the considered coalescence model). It also calculates the average fraction of the sampled genomes that were brought in by recombination (recombination coverage, *c*). mcorr was implemented with default settings and 1000 bootstraps.

Rates of recombination to mutation ($\delta/\mu$) were >1 for both *S. mitis* and *S. pneumoniae* (Supplementary Table 3), showing recombination to have had a greater role in the species diversification than mutation in both species and replicating previous results for *S. pneumoniae* [3, 4].

Mutational divergence ($\theta_{pool}$) is significantly higher than the sample's diversity for both species, indicating their population gene pool is highly diverse and that recombination within species can happen between highly divergent sequences (with divergences as high as 18% for *S. mitis* and 10% in *S. pneumoniae*). Although mutational divergence is similar between species (see also Supplementary Note 3), recombination divergence is considerably higher in *S. mitis* (Supplementary Table 3). Recombination frequency in both species was also evaluated by analysing the decay of LD with physical distance (Supplementary Fig. 3). The LD statistic $r^2$ between all pairs of synonymous SNVs with minor allele frequency bigger than 0.01 was calculated with PLINK 1.9 [5] up to a distance of 10 Kb (*--ld-window-k 10*), and using the command flags *--ld-window 5000* to allow computation of $r^2$ between all SNVs, and *--ld-window-r2 0* to obtain the tabling of all r2 values (>0).

Neutral LD $r^2$ values for *S. pneumoniae* decay from a maximum of ~0.3 to low, considered non-significant, values - less than 0.1 - within 1500bps; LD $r^2$ maximum value is ~ 0.06 for *S. mitis* and halves within 150 bps. These values together with mcorr recombination parameters, indicate a history of frequent genetic exchange that is preventing the presence of strong population structure in both species. The LD patterns for *S. mitis* are consistent with a quasi-sexual mode of evolution for this species [6].

Importantly, neutral LD $r^2$ values in *S. mitis* are considerably lower and decay to minimal values faster than LD in *S. pneumoniae*. In non-structured populations such as the ones under study, neutral LD between alleles is maintained by a balance between genetic drift and recombination [7]. One explanation for the observed differences in long-range LD for both species is that recombination rates are higher in *S. mitis*. However, under this hypothesis, recombination to mutation rates in serotype lineages in *S. pneumoniae* would be expected to be consistently lower than in *S. mitis*, and this pattern is not observed [4]. Therefore, the most likely explanation for lower neutral LD in *S. mitis* is long-term evolution based on lower levels of genetic drift. This hypothesis is also consistent with the genomic diversity and population structure patterns observed in *S. mitis* (see Discussion) and is compatible with its quasi-sexual mode of evolution described above.

#### Supplementary Note 2. Approximate Bayesian Computation

We further investigated the demographic history of *S. pneumoniae* under an approximated Bayesian computation (ABC) framework. The coalescent simulator FastSimBac [8] was used to generate 25,500 sequences under an exponential growth model, which incorporated mutation and recombination. Due to computational constrains, we simulated 100Kb segments with sample size and parameters matching the ones estimated for overlapping sliding windows of the same size across *S. pneumoniae* sampled genomes with an offset of 10 Kb between adjacent windows. The windows map to the first 930kb of the reference genome where read coverage is consistently high (>10% of mean depth of coverage) among all isolates (encompass a continuous genomic region of the core genome), and where the number of observed segregating sites in consistent across windows (Supplementary Dataset 5). Analysis of 150 Kb or 200 Kb windows led to similar genetic diversity (#seggregating sites/Kb) and Tajima's D values. In each simulation, parameters were randomly drawn from the following priors. Exponential growth rate was taken from an exponential distribution with rate 0.05. Recombination rate was taken from a gamma distribution with shape parameter 0.14*8 and rate parameter 8, truncated at a recombination rate of 0.3 (pre-truncation mean = 0.14, as estimated on observed data; Supplementary Note 1 and Supplementary Table 3). Mutation rate was set in each run by drawing a number of segregating sites *S* from a uniform distribution with boundaries selected at random from the observed number of segregating sites estimated in 100Kb-sized windows across *S. pneumoniae* sampled genomes, and scaling *S* with the mean number of segregating sites obtained from additional three FastSimBac runs with the same growth rate, recombination rate drawn from the prior described above, and a mutation rate of 1. Under a coalescence framework, the mean number of segregating sites obtained from these three simulations gives us a rough estimate of the expected length of the whole genealogy, which can then be used to properly scale the mutation rate. We only performed three simulations due to computational constrains, however we sought that these were sufficient to average out the variation in the independent estimates, since the number of segregating sites simulated across simulations was similar to the observed ones (Supplementary Fig. 4).

For every simulated and each of the 83 observed 100 kb segments, a set of summary statistics genetic diversity was recorded: the 5% percentiles of the minor allele frequencies; the 10% percentiles of pairwise Hamming distances between sequences, scaled by their maximum within the sample (simulated or observed); and Tajima's D. Calculations on observed data were based on synonymous (neutral) variation.

To estimate posterior distributions and median point estimates of the exponential growth rate, we used the Random Forest ABC model selection procedure (ABC-RF) from the *abcrf* R-package [9]. The normalized mean absolute error (NMAE) of the posterior parameter estimates (growth rates) obtained from the ABC-RF analysis was 0.15 (estimated via a built-in out-of-bag cross-validation), attesting to the accuracy of the random-forest classifier.

Despite only using synonymous variation, Tajima's D was observed to differ across the 83 windows in *S. pneumoniae*, leading to variable estimates of growth rate

(Supplementary Dataset 5). This variation is mostly driven by windows with genes found by *dN/dS* calculation to be under selection (*dN/dS*>2 or *dN/dS*<0.11; Supplementary Dataset 5). Focusing on the windows without such genes, we performed graphical posterior predictive checks by calculating the Tajima's D of simulated 1,500 100Kb-windows obtained using as parameters the 25% percentile, the median, and the 75% percentile of the estimated growth rates, and comparing these values with the observed Tajima's D of the corresponding windows from which the growth rates were extracted from (Supplementary Fig. 5). Graphical comparison allowed us to conclude that our approach recovers the observed Tajima's D well (Supplementary Fig. 5).

We have also performed a second set of 5600 simulations (dataset B; Supplementary Dataset 5) with the same population growth and mutation rate priors, but with recombination being drawn from a gamma distribution with shape parameter 0.14*40 and rate parameter 40, truncated at a recombination rate of 0.3 (same pre-truncation mean of 0.14). The NMAE of the posterior growth rates was 0.14. The resulting median parameter posterior estimates of this simulation set were on average differing only by 11% (at most by 34%) compared to the results of the first simulation setting. The median across all windows differs only by 7% from the estimate from the first simulation setup. This shows that the ABC analysis is robust towards a potential misspecification of the recombination rate (Supplementary Dataset 5).

Under this model, the median of the posterior growth rate distributions obtained across considered genomic windows is consistently > 1 for the two simulation sets (Supplementary Dataset 5). Our model shows then support for exponential growth in the *S. pneumoniae* population.

#### Supplementary Note 3. Higher neutral genetic diversity in *S. mitis* than in *S. pneumoniae* is not due to differences in their mutation rate

We confirmed experimentally and bioinformatically that the observed differences in neutral genetic diversity between *S. mitis* and *S. pneumoniae* are not due to differences in mutation rate between the species.

Firstly, we compared experimental determinations of spontaneous mutation rates for antibiotic resistant markers between the two species. Cells were grown to mid-exponential phase in liquid culture for 18 hours at 37°C, 5% $CO_2$. Three strains from the *S. mitis* dataset (one from each host population) were randomly selected and tested along with three laboratory pneumococcal strains (R6, G54, D39 or TIGR4) in biological triplicate. Minimum inhibitory concentrations of Rifampicin and Streptomycin were determined via 10-fold serial dilutions of antibiotic for each strain under investigation to ensure a suitably high level of antibiotic was used (i.e., that the bacterium did not have pre-existing resistance). Concentrated cultures (OD600 = 0.1) were plated on 32μg/ml Rifampicin and 500μg/ml Streptomycin in triplicate on BHI agar, 3% defibrinated horse blood (Oxoid) plates and grown for 24 hours at 37°C, 5% $CO_2$. Following incubation, discrete colonies were counted. For normalisation across strains, mutation rate (μ) was calculated per cell division from the mean number of (normally distributed) colonies counted across repeats. This calculation was under the assumptions that mutations arise throughout the cell cycle and cells are grown in an asynchronous population [10, 11]:

$$\mu = \frac{ln2m}{N_t - 1}$$

Where, m = mean number of mutations per culture (i.e., number of colonies)

$Nt$ = total number of starting cells

Resistance to both antibiotics can be accomplished through a singular point mutation. For both antibiotics, all *S. mitis* (and *S. pneumoniae*) strains generated discrete colonies, indicating occurrence of point mutations and a functional DNA mismatch repair system (as known for the *S. pneumoniae* strains used;). Spontaneous mutation rates between antibiotics and between strains did not differ significantly (Supplementary Fig. 6).

Secondly, we evaluated the effects of UV-exposure on cell viability in both species. Concentrated liquid cultures (OD600 = 0.1) of each of the three *S. mitis* and three *S. pneumococcus* strains were grown in 50ml BHI (Oxoid). Cultures were centrifuged at 4000rpm for 10 minutes to pellet cells, which were resuspended in 20ml 0.9% NaCl. Half of the resuspension (depth of resuspension in petri dish = 3.75mm) was exposed to a UV radiation source producing 10 J/min for 45 seconds. 10-fold serial dilutions (up to $10^{-8}$) of exposed and non-exposed culture were pipetted onto BHI, 3% horse blood agar plates in triplicate and incubated for 24 hours at 37°C, 5% $CO_2$ (De Ste Croix, 2017). Following incubation, discrete colonies were counted and colony forming units per ml (CFU/ml) were calculated.
After UV exposure, discrete colonies were identified across all six strains, and the number of CFU/ml was not significantly different between exposure/non exposure states. This confirms effective DNA nucleotide excision repair removing pyrimidine dimers generated by UVR exposure, which allows DNA replication to occur and to bacterial growth to ensue (Supplementary Fig 6B).

Thirdly, DNA repair during DNA replication was assessed by analysing the genetic variation of the *polA* gene, which encodes DNA polymerase I. This gene is highly variable, although a set of conserved residues have previously been proven essential for function [12]. All of these residues were ubiquitously present across isolates of both the *S. mitis* and *S. pneumoniae* datasets.

Finally, growth rate between species was experimentally compared. Liquid cultures were grown in BHI for 18 hours at 37°C, 5% CO2 and OD600 measurements were taken every hour for 24 hours.
An increased rate of DNA replication (faster growth) results in an accruement of genetic variation because of the inherent error rate of DNA replication (despite the proofreading capability of DNA polymerases). Thus, a higher rate of division accumulates more genetic variation. However, under laboratory conditions, the growth rate of *S. mitis* was comparable with *S. pneumoniae* (Supplementary Fig. 6C). Inherently, the doubling times were similar between the species, most likely reflecting similar generation times in natural conditions.

In combination, these results support *S. mitis* to generate de novo mutation at a comparable rate to that of *S. pneumoniae*, and therefore it cannot explain the differences in genetic diversity observed between species. Since it has been shown that spontaneous mutation rates for antibiotic resistance are consistently higher than mutation rates from mutation accumulation experiments with whole genome sequencing [13], we use a phylogenetic estimate determined for *S. pneumoniae* in our modelling of the population history of *S. mitis* and *S pneumoniae* [14]. A phylogenetic estimate of mutation rate allows to account for the generation time of these organisms in their natural environment.

#### Supplementary Tables

**Supplementary Table 1.** Pangenome statistics for *S. pneumoniae* and *S. mitis*. Pangenome and core genome sizes units are number of genes. N, sample size. Regional (continental) random sample of unrelated isolates show means of 1000 random samples with size equal to that of the smaller regional sample size within each species. 'All random sample of unrelated isolates' in *S. pneumoniae* shows the mean of 1000 random samples with size equal to the observed *S. mitis* unrelated sample size.

| Sample | N | Pangenome size | Core Genome size | Mean no. genes per sample | Median no. genes per sample |
|---|---|---|---|---|---|
| *S. pneumoniae* | | | | | |
| All | 802 | 27006 | 902 | 2004 | 1999 |
| African | 222 | 6898 | 1067 | 2022 | 2010 |
| Asian | 480 | 8179 | 982 | 194 | 1991 |
| European | 100 | 5188 | 1169 | 2016 | 2020 |
| All unrelated | 353 | 8811 | 934 | 2005 | 1995 |
| African unrelated | 78 | 6282 | 1075 | 2035 | 2014 |
| Asian unrelated | 207 | 7192 | 1013 | 1991 | 1985 |
| European unrelated | 68 | 5047 | 1175 | 2014 | 2020 |
| African random sample of unrelated isolates | 68 | 6215 | 1080 | 2038 | 2015 |
| Asian random sample of unrelated isolates | 68 | 5818 | 1101 | 1983 | 1982 |
| All random sample of unrelated isolates | 75 | 6064 | 1092 | 2005 | 1995 |
| *S. mitis* | | | | | |
| All | 119 | 12005 | 948 | 1876 | 1828 |
| African | 32 | 5520 | 1058 | 1774 | 1772 |
| Asian | 36 | 5288 | 1091 | 1841 | 1835 |
| European | 49 | 7004 | 992 | 1884 | 1864 |
| All unrelated | 75 | 9626 | 951 | 1832 | 1816 |
| African unrelated | 25 | 5495 | 1058 | 1775 | 1773 |
| Asian unrelated | 18 | 5242 | 1093 | 1847 | 1834 |
| European unrelated | 32 | 6897 | 993 | 1869 | 1862 |
| African random sample of unrelated isolates | 18 | 4764 | 1082 | 1761 | 1772 |
| European random sample of unrelated isolates | 18 | 5322 | 1071 | 1869 | 1864 |

**Supplementary Table 2**. Between population divergence estimates (Hudson's FST +/- SE). *S. mitis*, above diagonal; *S. pneumoniae*, below diagonal.

| | Africa | Asia | Europe |
|---|---|---|---|
| Africa | | 0.0684 +/- 0.0017 | 0.0495 +/- 0.0011 |
| Asia | 0.0306 +/- 0.0035 | | 0.0107 +/- 0.0012 |
| Europe | 0.0406 +/- 0.0027 | 0.0464 +/- 0.0038 | |

**Supplementary Table 3**. mcorr estimates of recombination and mutation parameters for *S. mitis* and *S. pneumoniae*. n, sample size; $n_{genes}$, number of genes analysed; $d_{sample}$, diversity of the sample; $\theta_{pool}$, mutational divergence; and $\phi_{pool}$, recombinational divergence of the of the species' gene pool; $\delta/\mu$, the relative rate of recombination to mutation; $\bar{f}$, the mean recombination fragment length; *c*, recombination coverage. A definition of these parameters is in material and methods.

| Species | n | $n_{genes}$ | $d_{sample}$ | $\theta_{pool}$ | $\phi_{pool}$ | $\delta/\mu$ | $\bar{f}$ (bp) | *c* |
|---|---|---|---|---|---|---|---|---|
| *S. mitis* | 75 | 950 | 0.046 | 0.18 | 0.82 | 4.61 | 2511 | 0.32 |
| *S. pneumoniae* | 353 | 972 | 0.012 | 0.10 | 0.14 | 1.41 | 1004 | 0.13 |

**Supplementary Table 4**. Genome accession and geographical origin of *Streptococcus mitis* genomes.

| Host | isolate | region | Biosample | Bioproject |
|---|---|---|---|---|
| B1015B24 | B1015B24C5 | Africa | SAMEA8000357 | PRJEB42963 |
| B1015G24 | B1015G24C3 | Africa | SAMEA8000358 | PRJEB42963 |
| B1015H24 | B1015H24C1 | Africa | SAMEA8000359 | PRJEB42963 |
| B1015H24 | B1015H24C3 | Africa | SAMEA8000360 | PRJEB42963 |
| B1015H24 | B1015H24C4 | Africa | SAMEA8000361 | PRJEB42963 |
| S1071B24 | S1071B24C1 | Africa | SAMEA8000362 | PRJEB42963 |
| S1071B24 | S1071B24C2 | Africa | SAMEA8000363 | PRJEB42963 |
| S1071G24 | S1071G24C1 | Africa | SAMEA8000364 | PRJEB42963 |
| S1071G24 | S1071G24C2 | Africa | SAMEA8000365 | PRJEB42963 |
| S1071G24 | S1071G24C3 | Africa | SAMEA8000366 | PRJEB42963 |
| S1071G24 | S1071G24C4 | Africa | SAMEA8000367 | PRJEB42963 |
| S1071G24 | S1071G24C5 | Africa | SAMEA8000368 | PRJEB42963 |
| S1072B24 | S1072B24C4 | Africa | SAMEA8000369 | PRJEB42963 |
| S1072B24 | S1072B24C5 | Africa | SAMEA8000370 | PRJEB42963 |
| S1072G24 | S1072G24C4 | Africa | SAMEA8000371 | PRJEB42963 |
| S1075M24 | S1075M24C2 | Africa | SAMEA8000372 | PRJEB42963 |
| S1075M24 | S1075M24C3 | Africa | SAMEA8000373 | PRJEB42963 |
| S1075M24 | S1075M24C5 | Africa | SAMEA8000374 | PRJEB42963 |
| S1086G24 | S1086G24C1 | Africa | SAMEA8000375 | PRJEB42963 |
| S1088B24 | S1088B24C2 | Africa | SAMEA8000376 | PRJEB42963 |
| S1088M24 | S1088M24C1 | Africa | SAMEA8000377 | PRJEB42963 |
| S1091B24 | S1091B24C4 | Africa | SAMEA8000378 | PRJEB42963 |
| S1092G24 | S1092G24C1 | Africa | SAMEA8000379 | PRJEB42963 |
| S1092G24 | S1092G24C3 | Africa | SAMEA8000380 | PRJEB42963 |
| S1092G24 | S1092G24C4 | Africa | SAMEA8000381 | PRJEB42963 |
| S1093G24 | S1093G24C1 | Africa | SAMEA8000382 | PRJEB42963 |
| S1093G24 | S1093G24C2 | Africa | SAMEA8000383 | PRJEB42963 |
| S1093G24 | S1093G24C3 | Africa | SAMEA8000384 | PRJEB42963 |
| S1096G24 | S1096G24C5 | Africa | SAMEA8000385 | PRJEB42963 |
| S1096M24 | S1096M24C3 | Africa | SAMEA8000386 | PRJEB42963 |
| S1096M24 | S1096M24C5 | Africa | SAMEA8000387 | PRJEB42963 |
| C1 | C1C1 | Asia | SAMN37997237 | PRJNA1032561 |
| C1 | C1C2 | Asia | SAMN37997238 | PRJNA1032561 |
| C1 | C1C3 | Asia | SAMN37997239 | PRJNA1032561 |
| C1 | C1C4 | Asia | SAMN37997240 | PRJNA1032561 |
| C1 | C1C5 | Asia | SAMN37997241 | PRJNA1032561 |
| C1 | C1T1 | Asia | SAMN37997242 | PRJNA1032561 |

| | | | | |
|---|---|---|---|---|
| C1 | C1T3 | Asia | SAMN37997243 | PRJNA1032561 |
| C2 | C2C1 | Asia | SAMN37997244 | PRJNA1032561 |
| C2 | C2C2 | Asia | SAMN37997245 | PRJNA1032561 |
| C2 | C2C3 | Asia | SAMN37997246 | PRJNA1032561 |
| C2 | C2C4 | Asia | SAMN37997247 | PRJNA1032561 |
| C2 | C2C5 | Asia | SAMN37997248 | PRJNA1032561 |
| C2 | C2C6 | Asia | SAMN37997249 | PRJNA1032561 |
| C2 | C2T1 | Asia | SAMN37997250 | PRJNA1032561 |
| C2 | C2T2 | Asia | SAMN37997251 | PRJNA1032561 |
| C2 | C2T3 | Asia | SAMN37997252 | PRJNA1032561 |
| C2 | C2T4 | Asia | SAMN37997253 | PRJNA1032561 |
| C3 | C3C1 | Asia | SAMN37997254 | PRJNA1032561 |
| C3 | C3C2 | Asia | SAMN37997255 | PRJNA1032561 |
| C3 | C3C3 | Asia | SAMN37997256 | PRJNA1032561 |
| C3 | C3C4 | Asia | SAMN37997257 | PRJNA1032561 |
| C3 | C3T1 | Asia | SAMN37997258 | PRJNA1032561 |
| C4 | C4C1 | Asia | SAMN37997259 | PRJNA1032561 |
| C4 | C4C2 | Asia | SAMN37997260 | PRJNA1032561 |
| C4 | C4C3 | Asia | SAMN37997261 | PRJNA1032561 |
| C4 | C4C4 | Asia | SAMN37997262 | PRJNA1032561 |
| C5 | C5C1 | Asia | SAMN37997263 | PRJNA1032561 |
| C5 | C5C2 | Asia | SAMN37997264 | PRJNA1032561 |
| C5 | C5C3 | Asia | SAMN37997265 | PRJNA1032561 |
| C5 | C5C4 | Asia | SAMN37997266 | PRJNA1032561 |
| C5 | C5C5 | Asia | SAMN37997267 | PRJNA1032561 |
| C5 | C5T1 | Asia | SAMN37997268 | PRJNA1032561 |
| C5 | C5T2 | Asia | SAMN37997269 | PRJNA1032561 |
| C5 | C5T3 | Asia | SAMN37997270 | PRJNA1032561 |
| C5 | C5T4 | Asia | SAMN37997271 | PRJNA1032561 |
| C5 | C5T5 | Asia | SAMN37997272 | PRJNA1032561 |
| SK1126 | SK1126 | Asia | SAMN02836935 | PRJNA242574 |
| B1 | B1C1 | Europe | SAMN37997273 | PRJNA1032561 |
| B1 | B1C2 | Europe | SAMN37997274 | PRJNA1032561 |
| B1 | B1C3 | Europe | SAMN37997275 | PRJNA1032561 |
| B1 | B1C4 | Europe | SAMN37997276 | PRJNA1032561 |

| | | | | |
|---|---|---|---|---|
| B1 | B1C5 | Europe | SAMN37997277 | PRJNA1032561 |
| B1 | B1T1 | Europe | SAMN37997278 | PRJNA1032561 |
| B1 | B1T2 | Europe | SAMN37997279 | PRJNA1032561 |
| B2 | B2C1 | Europe | SAMN37997280 | PRJNA1032561 |
| B2 | B2C2 | Europe | SAMN37997281 | PRJNA1032561 |
| B2 | B2C3 | Europe | SAMN37997282 | PRJNA1032561 |
| B2 | B2C4 | Europe | SAMN37997283 | PRJNA1032561 |
| B2 | B2T1 | Europe | SAMN37997284 | PRJNA1032561 |
| B2 | B2T2 | Europe | SAMN37997285 | PRJNA1032561 |
| B2 | B2T3 | Europe | SAMN37997286 | PRJNA1032561 |
| B2 | B2T4 | Europe | SAMN37997287 | PRJNA1032561 |
| B3 | B3C1 | Europe | SAMN37997288 | PRJNA1032561 |
| B3 | B3C3 | Europe | SAMN37997289 | PRJNA1032561 |
| B3 | B3C4 | Europe | SAMN37997290 | PRJNA1032561 |
| B3 | B3T1 | Europe | SAMN37997291 | PRJNA1032561 |
| B3 | B3T2 | Europe | SAMN37997292 | PRJNA1032561 |
| B4 | B4C1 | Europe | SAMN37997293 | PRJNA1032561 |
| B4 | B4C2 | Europe | SAMN37997294 | PRJNA1032561 |
| B4 | B4C3 | Europe | SAMN37997295 | PRJNA1032561 |
| B4 | B4C4 | Europe | SAMN37997296 | PRJNA1032561 |
| B4 | B4T1 | Europe | SAMN37997297 | PRJNA1032561 |
| B5 | B5C1 | Europe | SAMN37997298 | PRJNA1032561 |
| B5 | B5C2 | Europe | SAMN37997299 | PRJNA1032561 |
| B5 | B5C3 | Europe | SAMN37997300 | PRJNA1032561 |
| B5 | B5C4 | Europe | SAMN37997301 | PRJNA1032561 |
| B5 | B5C5 | Europe | SAMN37997302 | PRJNA1032561 |
| B5 | B5T1 | Europe | SAMN37997303 | PRJNA1032561 |
| B5 | B5T2 | Europe | SAMN37997304 | PRJNA1032561 |
| SK1073 | SK1073 | Europe | SAMN00621702 | PRJNA66111 |
| SK1080 | SK1080 | Europe | SAMN00621705 | PRJNA66113 |
| SK137 | SK137 | Europe | SAMN03334900 | PRJNA274768 |
| SK145 | SK145 | Europe | SAMN03334902 | PRJNA274768 |
| SK271 | SK271 | Europe | SAMN02698681 | PRJNA242568 |
| SK321 | SK321 | Europe | SAMN00001433 | PRJNA33353 |
| SK564 | SK564 | Europe | SAMN00001434 | PRJNA33355 |
| SK569 | SK569 | Europe | SAMN00621701 | PRJNA67187 |
| SK575 | SK575 | Europe | SAMN00761799 | PRJNA75135 |

| | | | | |
|---|---|---|---|---|
| SK578 | SK578 | Europe | SAMN02836940 | PRJNA242567 |
| SK579 | SK579 | Europe | SAMN00761835 | PRJNA75157 |
| SK597 | SK597 | Europe | SAMN00001435 | PRJNA33357 |
| SK608 | SK608 | Europe | SAMN02836941 | PRJNA242555 |
| SK616 | SK616 | Europe | SAMN00761793 | PRJNA75129 |
| SK629 | SK629 | Europe | SAMN02836936 | PRJNA242570 |
| SK637 | SK637 | Europe | SAMN02836939 | PRJNA242571 |
| SK642 | SK642 | Europe | SAMN02836938 | PRJNA242572 |
| SK667 | SK667 | Europe | SAMN02836937 | PRJNA242573 |

### Supplementary Figures

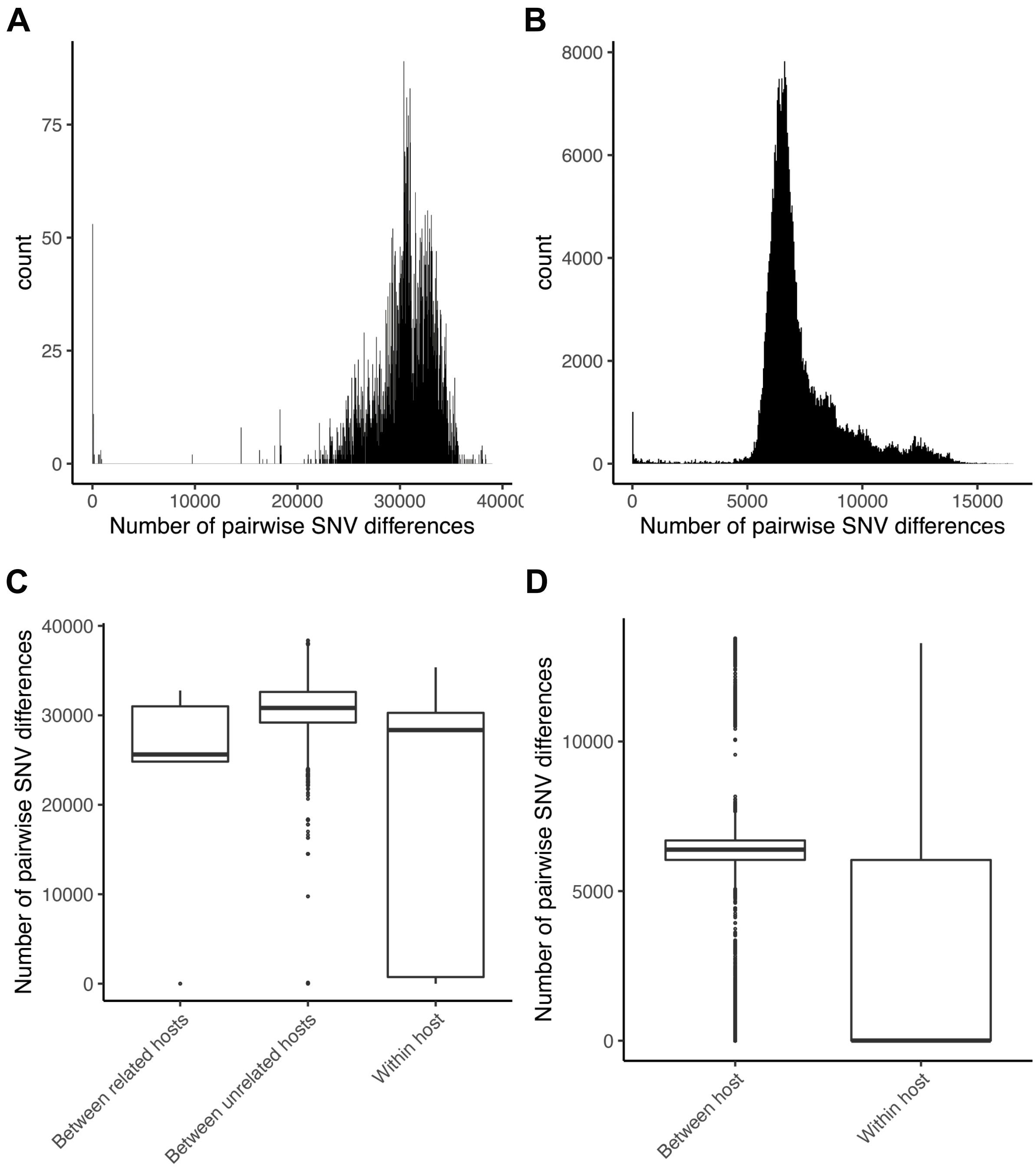


**Supplementary Figure 1.** Pairwise SNV differences distribution for *S. mitis* and *S. pneumoniae*. **A** Pairwise SNV differences distribution for *S. mitis* total sample (n=119). **B** Pairwise SNV differences distribution for *S. pneumoniae* total sample (n=810). **C** Pairwise SNV differences distribution between related hosts, between unrelated hosts and within host for *S. mitis*. **D** Pairwise SNV differences distribution between unrelated hosts and within hosts for *S. pneumoniae* (for the African dataset, n=230).

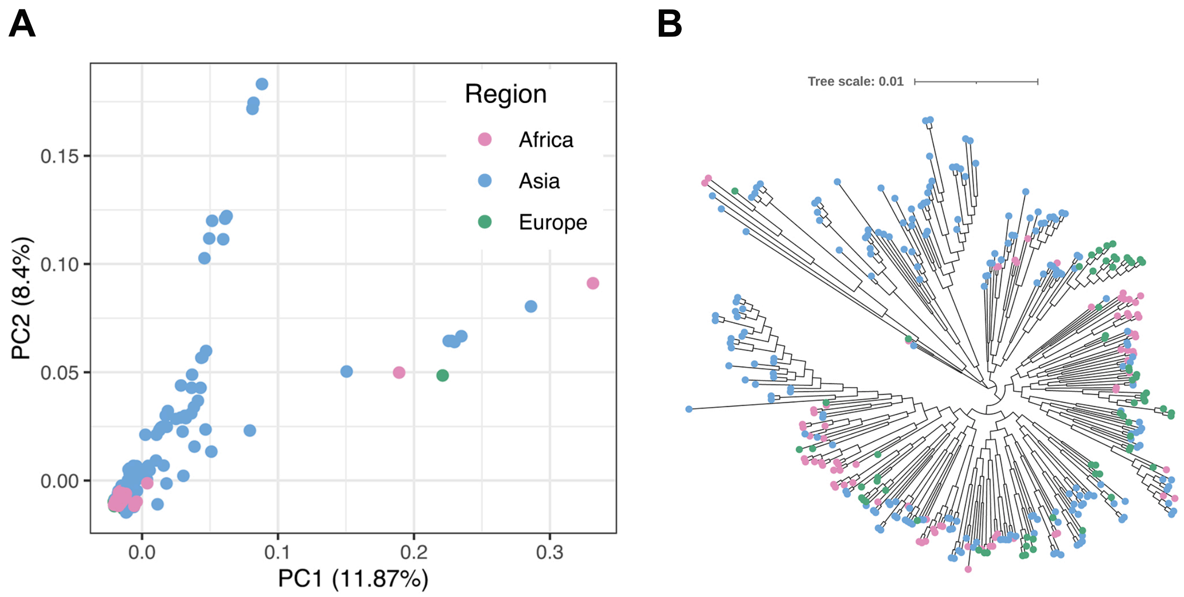


**Supplementary Figure 2.** Phylogenetic and population structure for *S. pneumoniae*'s total sample (including serotype NT). **A** PCA computed in Plink v1.9 [5]. **B** Maximum likelihood unrooted phylogenetic tree obtained using FastTree [15]. There are subclades of Asian lineages which belong exclusively to the PCA outlying serotypes NT (three subclades with isolates belonging mainly to GPSCs 28, 42, 60, 66, 118) and 19F (one subclade classified as GPSC-1), although both NT and 19F clades also include African and European lineages. The NT cluster (and specifically the Asian isolates) comprise nonencapsulated isolates which were previously reported to have higher recombination rates generating significantly more diversity within this cluster [16]. Colour code corresponds to geographic region: Africa, pink; Asia, blue; Europe, green.

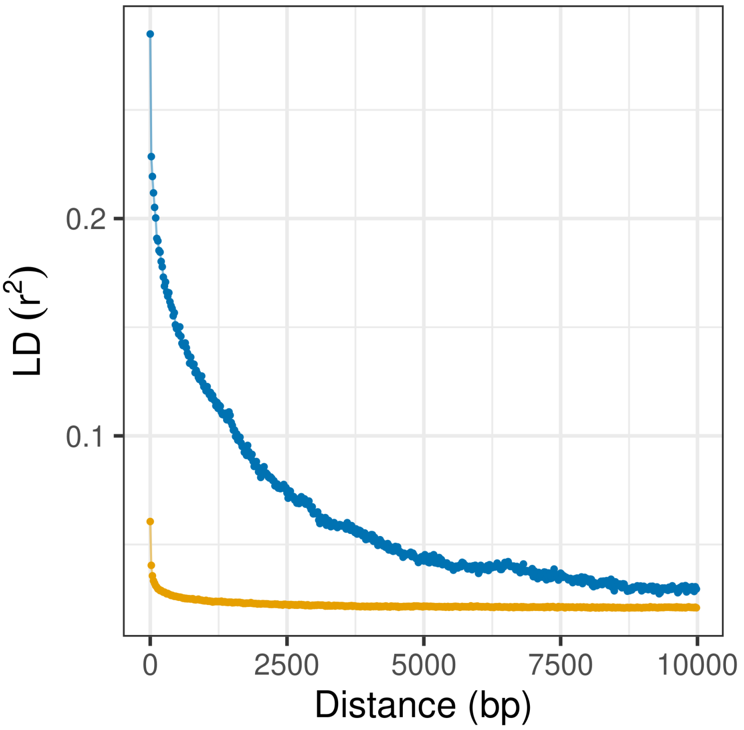


**Supplementary Figure 3.** LD decay with distance for *S. mitis* (orange) and *S. pneumoniae*(blue).

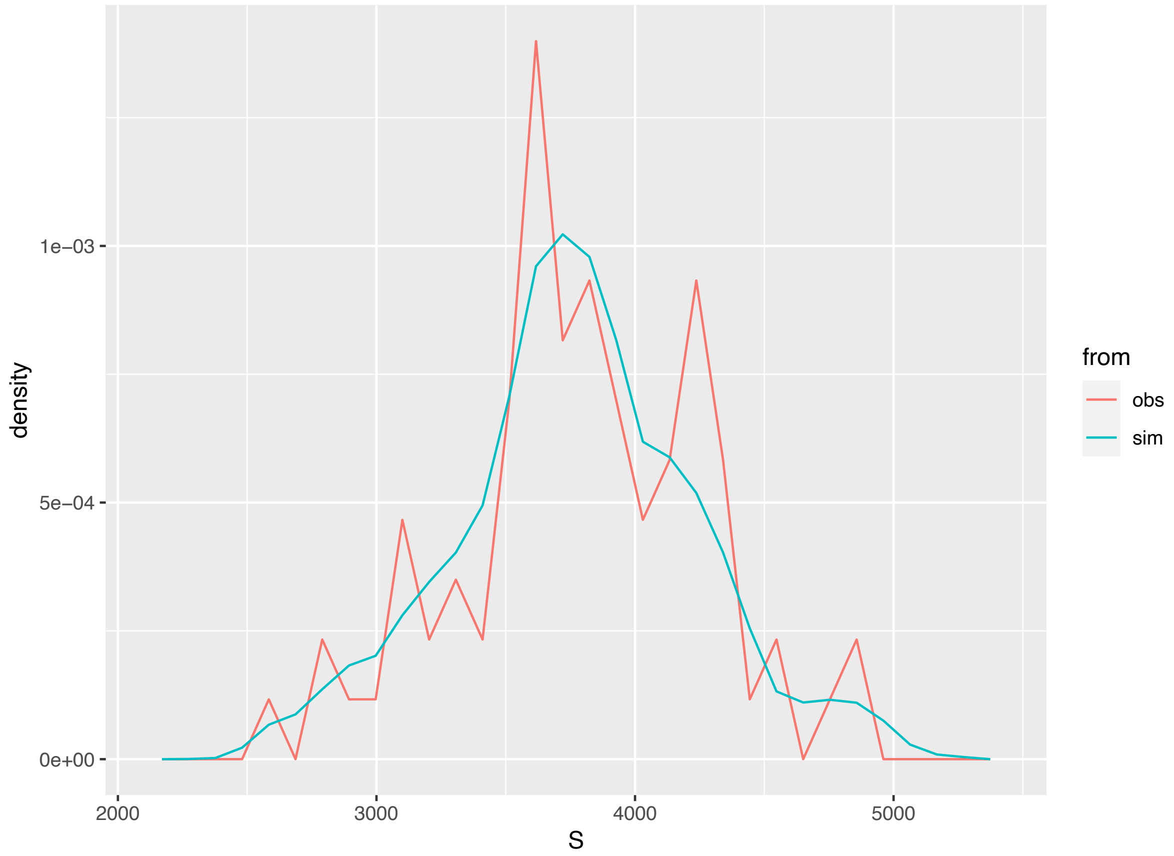


**Supplementary Figure 4.** Comparison between the distribution of observed number of segregating sites (pink; across 83 *S. pneumoniae* core genome windows) and the distribution of simulated number of segregating sites used in the ABC approach implemented to investigate *S. pneumoniae* demographic history. The distribution of simulated S values follows the observed distribution closely. S, number of segregating sites.

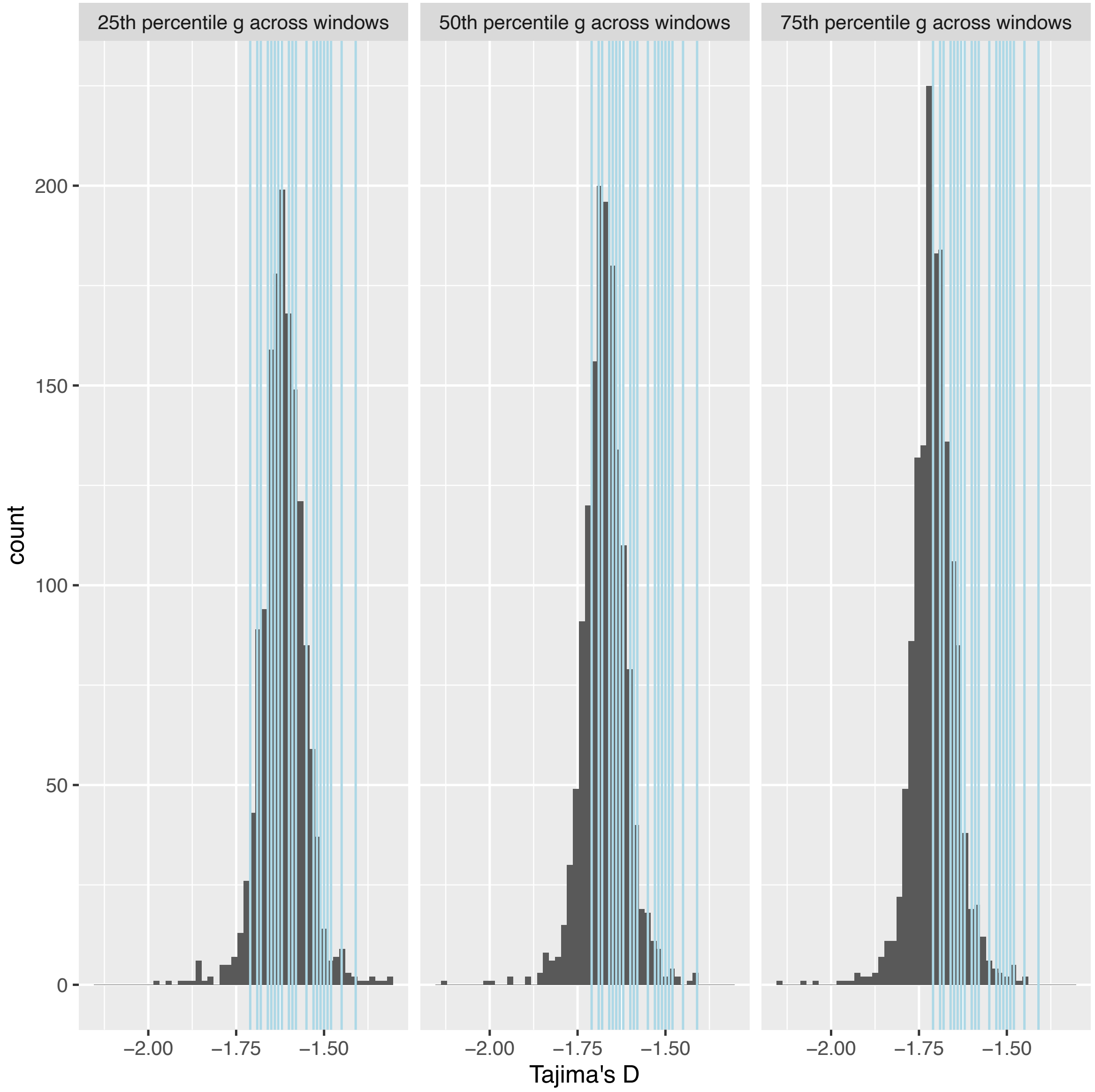


**Supplementary Figure 5.** Posterior predictive checks of the ABC approach implemented to investigate *S. pneumoniae* demographic history. The histograms show, from left to right, the distribution of Tajima's D obtained from simulating 1,500 100Kb-windows using as parameters the 25% percentile, the median, and the 75% percentile of estimated growth rates from across windows without genes under selection (Supplementary Data 5). The vertical blue bars correspond to the observed Tajima's D estimated across the 26 windows considered.

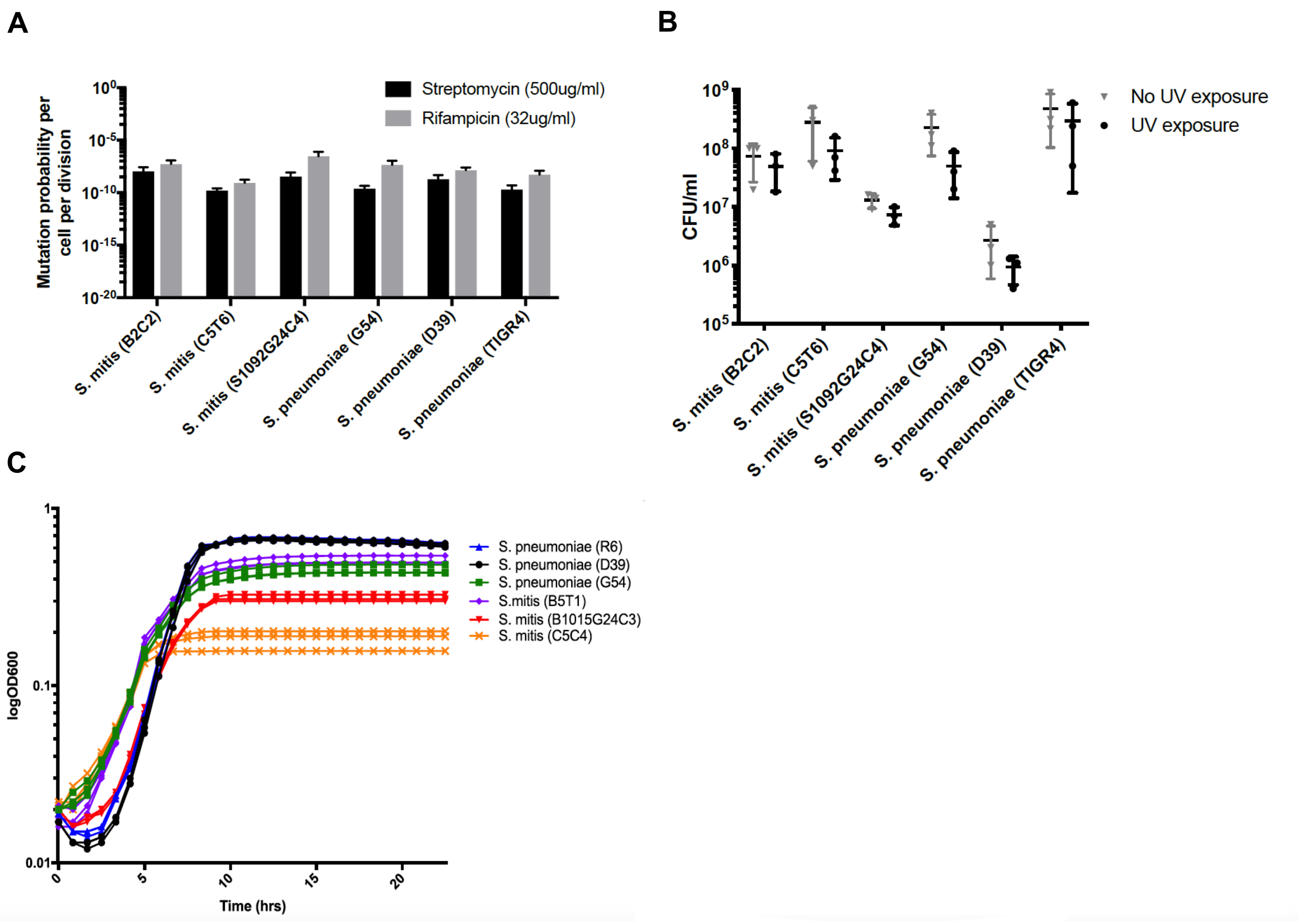


**Supplementary Figure 6.** Experimental confirmation of comparable mutation rates between *S. mitis* and *S. pneumoniae***.** Three isolates per species per experiment were used, which were performed in biological triplicate. Error bars show SD from mean. **A** Spontaneous mutation rate for inhibitory concentration of Streptomycin (black) and Rifampicin (grey). No significant difference between species was identified (two-way ANOVA) for either Streptomycin (*P-value* = 0.36) or Rifampicin (*P-value* = 0.41). **B** Cell viability with and without UV exposure. Number of viable cells was not statistically significant at the 0.05 level between cells exposed to UV and those that were not. Significance between states tested by unpaired t-test (*P-value*: B2C2=0.38, C5T6=0.36, S1092G24C4=0.09, G54=0.08, D39=0.69, TIGR4=0.43). **C** Twenty-four-hour growth curves from starting OD600 of 0.002 (reaching 0.157-0.642) demonstrating comparable growth rate between species.

13. Lee H, Popodi E, Tang H, Foster PL. Rate and molecular spectrum of spontaneous mutations in the bacterium Escherichia coli as determined by whole-genome sequencing. Proc

Natl Acad Sci U S A. 2012;109(41):E2774-83. Epub 20120918. doi: 10.1073/pnas.1210309109. PMID: 22991466; PMCID: PMCPMC3478608.

14. Croucher NJ, Harris SR, Fraser C, Quail MA, Burton J, van der Linden M, et al. Rapid pneumococcal evolution in response to clinical interventions. Science. 2011;331(6016):430-4. doi: 10.1126/science.1198545. PMID: 21273480; PMCID: PMCPMC3648787.

15. Price MN, Dehal PS, Arkin AP. FastTree 2-Approximately Maximum-Likelihood Trees for Large Alignments. Plos One. 2010;5(3). doi: 10.1371/journal.pone.0009490. PMID: WOS:000275328800002.

16. Chewapreecha C, Harris SR, Croucher NJ, Turner C, Marttinen P, Cheng L, et al. Dense genomic sampling identifies highways of pneumococcal recombination. Nat Genet. 2014;46(3):305-+. doi: 10.1038/ng.2895. PMID: WOS:000332036700016.
